# Comparison of Single-cell Long-read and Short-read Transcriptome Sequencing of Patient-derived Organoid Cells of ccRCC: Quality Evaluation of the MAS-ISO-seq Approach

**DOI:** 10.1101/2024.03.14.584953

**Authors:** Natalia Zajac, Qin Zhang, Anna Bratus-Neuschwander, Weihong Qi, Hella Anna Bolck, Tülay Karakulak, Tamara Carrasco Oltra, Holger Moch, Abdullah Kahraman, Hubert Rehrauer

## Abstract

Single-cell RNA sequencing is used in profiling gene expression differences between cells. Short-read sequencing platforms provide high throughput and high-quality information at the gene-level, but the technique is hindered by limited read length, failing in providing an understanding of the cell heterogeneity at the isoform level. This gap has recently been addressed by the long-read sequencing platforms that provide the opportunity to preserve full-length transcript information during sequencing. To objectively evaluate the information obtained from both methods, we sequenced four samples of patient-derived organoid cells of clear cell renal cell carcinoma and one healthy sample of kidney organoid cells on Illumina Novaseq 6000 and PacBio Sequel IIe. For both methods, for each sample, the cDNA was derived from the same 10x Genomics 3’ single-cell gene expression cDNA library. Here we present the technical characteristics of both datasets and compare cell metrics and gene-level information. We show that the two methods largely overlap in the results but we also identify sources of variability which present a set of advantages and disadvantages to both methods.

## Introduction

Short-read sequencing has been the basis for single-cell RNA sequencing, a method for gene expression profiling at a resolution of single cells (Dong et al., 2021; Heumos et al., 2023). Rapid developments in library protocols in the field have allowed collecting data from multiple modalities from a single cell, including chromatin accessibility (ATAC-seq) (Buenrostro et al., 2015; Cusanovich et al., 2015; Heumos et al., 2023), surface protein abundance (CITE-seq) (Hao et al., 2021; Heumos et al., 2023; Stoeckius et al., 2017), adaptive immune receptor repertoire (Pai & Satpathy, 2021), nucleosome occupancy (Clark et al., 2018; Hao et al., 2021; Pott, 2017) or spatial information (Vickovic et al., 2019).

However, due to transcript fragmentation necessary for short-read sequencing, the data does not provide isoform-level information or the information on structural variants, which can have a profound impact on deciphering cell-specific heterogeneity (Dondi et al., 2023; Hård et al., 2023; Pardo-Palacios et al., 2023). The current options of studying isoforms with short reads is limited to their partial reconstruction (SMART-Seq3) (Hagemann-Jensen et al., 2020) or through the use of probabilistic methods (Pan et al., 2022). In consequence, the community has turned to long-read sequencing that allows for full-length transcript sequencing (Pardo-Palacios et al., 2023). Historically, the long-read data has often been coupled with short-read data due to lacking in coverage (mostly concerning Pacific Biosciences, PacBio, data) or due to lower read accuracy (mostly concerning Oxford Nanopore data) (Dong et al., 2021; Tian et al., 2021). Recent developments in both technologies including improvements in read accuracy or throughput and offering of novel solutions, such as direct RNA sequencing (Depledge et al., 2019; Jain et al., 2022), have allowed the decoupling of long- and short-read approaches. The Multiplexed Array Sequencing method (MAS-seq) offered by PacBio, released in October 2022 and now relabelled as Kinnex, has offered higher throughput through concatenating full-length transcripts into longer fragments that can then be sequenced on one of their instruments (Al’Khafaji et al., 2021; PacBio, 2023a). Each fragment thus consists of an average of fifteen transcripts instead of one (PacBio, 2023b). After sequencing the reads can then be bioinformatically broken down to the original transcripts.

For those with interest in isoform information, the use of long reads without short reads can sound very promising, due to saving on costs and the amount of work, but poses a question of whether the data is comparable in the information provided by the short reads. Dondi et al. (2023) have partially addressed the question by comparing the expressed genes captured by both methods and cell type identification in ovarian cancer cells, but the research focused more on the additional information provided by long reads in isoform expression heterogeneity in tumour cells. In this study, we sequenced four patient-derived organoid cells of clear cell renal cell carcinoma (ccRCC) and one patient-derived metastasis-free kidney organoid cells using Illumina NovaSeq 6000, a short-read sequencing platform, and PacBio Sequel IIe, a long-read sequencing platform. We compare the datasets focusing on raw data characteristics (mapping rate, types of data artefacts, etc.). In particular, we make use of the fact that the protocol tags the cDNA molecules with cell barcodes and unique molecular identifiers (UMIs). This allows us to perform a per-molecule matched comparison of the short and long reads generated from the same original cDNA. Additionally, we evaluate the similarities and differences in computing the gene count matrix, and in consequence also the gene-level information obtained by both methods.

## Results

### Mapping of the reads to the genome

We generated short- and long-read sequencing data for four patient-derived organoid samples of ccRCC complimented with a matched sample of healthy kidney organoid cells. For both sequencing methods we used the same cDNA generated with 10x Genomics single-cell 3’ reagent kits (Fig 1.a). For short-read sequencing the library further underwent fragmentation and size selection and was then sequenced on NovaSeq 6000, therefore we refer to the data as “Illumina data” throughout the manuscript (Fig 1.a). For long-read sequencing we subsequently followed the MAS-Seq protocol and sequenced the data on Sequel IIe (Fig 1.a). We refer to the data as “PacBio data”. The data was purposefully targeted for a low number of cells (approx. 700) for deeper coverage per cell. With a total of 193-271 million reads per sample from the Illumina data, we obtained between 549 (ccRCC_2) to 1,664 (ccRCC_4) cells with 442,530 and 141,887 mean reads per cell, respectively (Supplementary Table 1). For PacBio data, we obtained between 29.4 and 58.3 million reads (generated from 1.88 to 3.79 HiFi reads of the concatenated constructs). We obtained between 310 (ccRCC_3) and 1,091 (ccRCC_4) cells per sample and the average number of reads per cell ranged from 21,499 (ccRCC_4) to 96,620 (ccRCC_2) (Supplementary Table 1).

**Figure 1.**
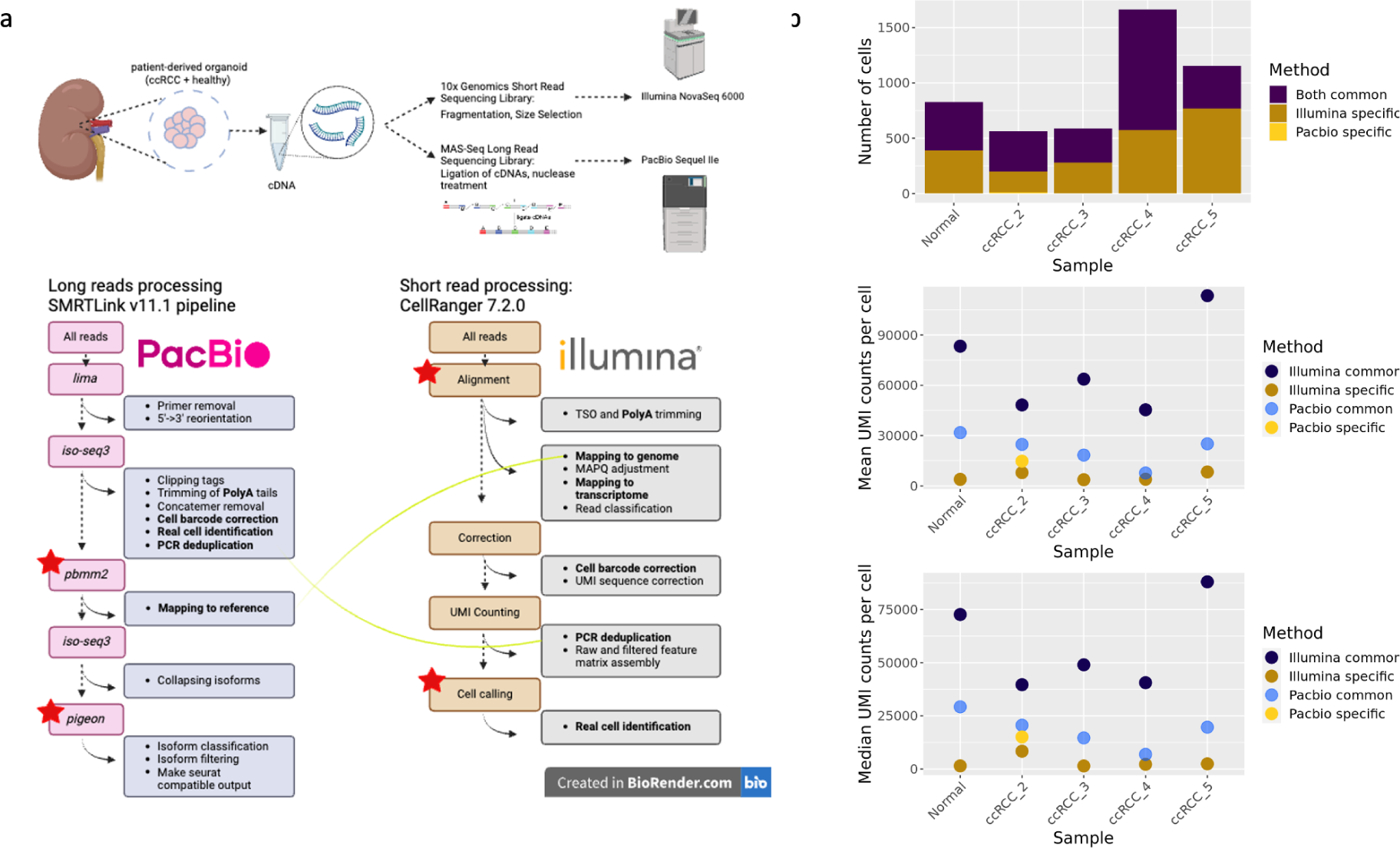
a) The scRNA-seq workflow: four samples of patient-derived organoid cells of ccRCC and one sample of healthy kidney organoid cells were used to generate 10x Genomics single-cell 3’ gene expression library that was then sequenced on two platforms generating long (PacBio) and short (Illumina) reads. The image of read concatenation performed during MAS-seq was derived from (Webber et al., 2023). The two datasets were then processed with state-of-the-art workflows. Long reads were analysed with SMRTLink v11.1 pipeline and short reads were processed with CellRanger v7.2.0. The common steps in the workflows are highlighted in bold. Key steps are joined with a yellow line (mapping and PCR duplicate removal). Red stars indicate the steps we compared in our analysis. b) Summary statistics (from top to bottom): the number of cells per sample identified in the two datasets, the mean and median UMI counts per cell for common cells and cells unique to each method.

All of the cells detected by PacBio were also found in Illumina (except for ccRCC_2 for which 12 cells were unique to PacBio) but CellRanger detected more cells in Illumina data for all samples (Fig 1.b). The reason stemmed from the difference in cell calling by the different processing algorithms. Iso-Seq3 by default uses the Knee Finding method to determine real cells which takes into account the number of UMIs per cell. CellRanger additionally uses the RNA profile and retains those barcodes with UMI counts below the threshold but whose RNA profile strongly disagrees with the background model (Lun et al., 2019). For Illumina-specific cells we observed a lower number of mean and median UMIs per cell (Fig 1.b). The highest mean and median UMIs per cell were observed for Illumina cells common with PacBio (Fig 1.b).

The Illumina raw data and the PacBio raw data were mapped to the same human reference genome (GrCh38.p13) and annotated with the same GENCODE 39 annotation. Prior to mapping, PacBio reads underwent PCR deduplication via clustering by UMI and cell barcodes. The step reduced the number of reads per sample to between 15.85 (ccRCC_3) and 27.8 million reads (Normal). The mapping rate for PacBio for all samples was then above 99.4%. For Illumina, PCR deduplication occurs only after mapping taking the read coordinates into account. Thus, 85.4% to 94% of reads mapped to the genome and between 73% and 85.6% of the reads mapped confidently (Supplementary Table 1).

Since the same cDNA has been sequenced with both methods, we could compare the Illumina and PacBio readouts for the same original cDNA molecules, with the same cell barcodes and UMIs. For these per-molecule comparisons we sampled approximately 5% of the cells. For Illumina data the number of unique cell barcode-UMI combinations (tag IDs) ranged from 1.3 million to 4.015 million with an average of 83,230-154,791 UMIs per cell. For PacBio data, the number of unique tag IDs ranged from 0.788 million to 1.07 million with an average of 43,791-60,978 UMIs per cell (Supplementary Table 2). Short-read sequencing clearly provided a higher coverage (Fig 2.a). The common tag IDs ranged from 595,022 to 896,729, leaving only 10-32.1% of tag IDs unique to PacBio but as much as 37.4-84.1% of tag IDs unique to Illumina (Fig 2.b).

**Figure 2.**
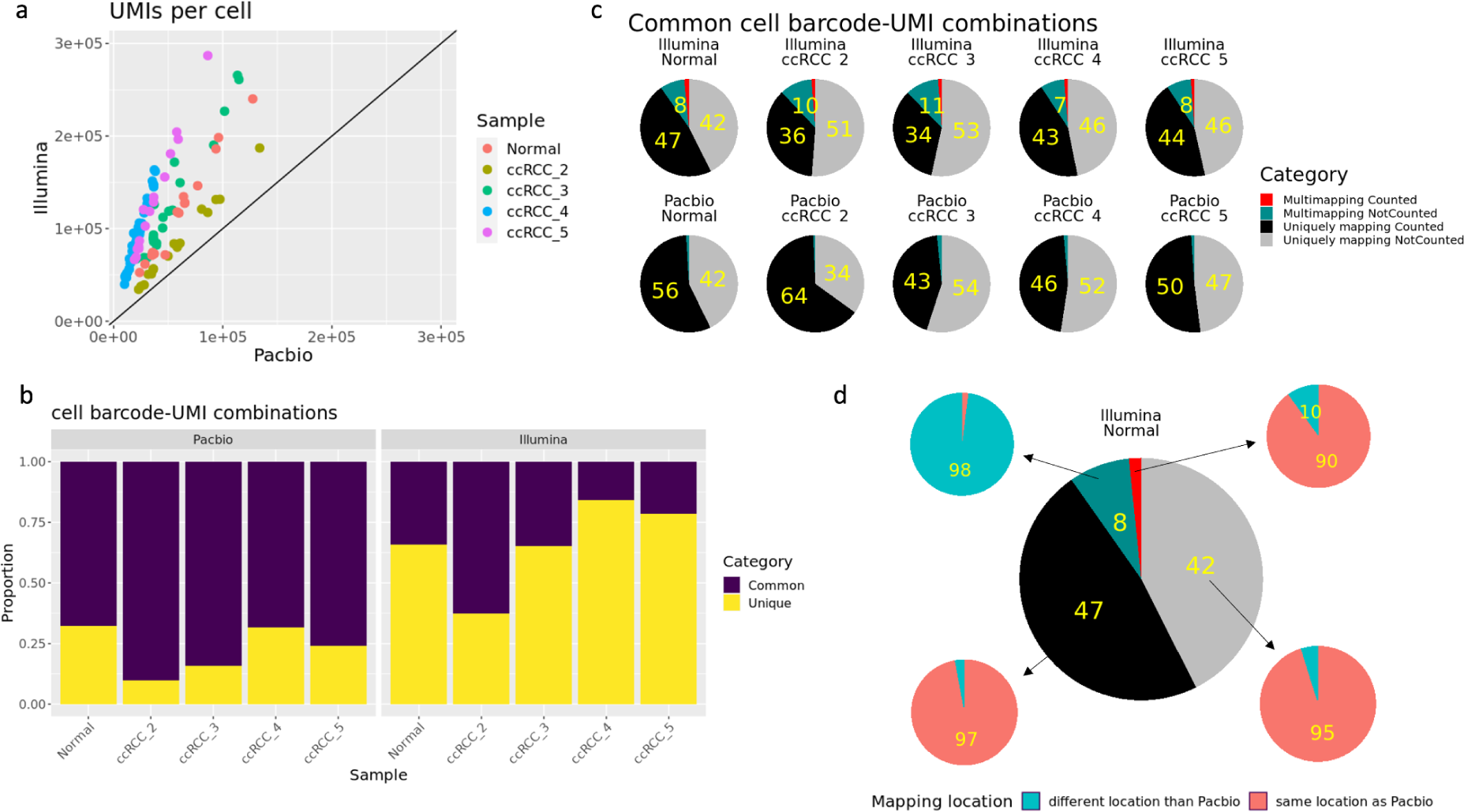
Comparison of matched Illumina and PacBio cell barcode-UMI combinations. a) Number of UMIs per cell for the subsampled cell barcodes. b) The proportion of cell barcode-UMI combinations common between the two datasets and unique to each dataset. c) The proportion of shared cell barcode-UMI combinations divided by alignment type (uniquely or multi-mapping) and by whether they were counted into the final gene count matrix (if below 5%, proportion not printed) d) The proportion of Illumina cell barcode-UMI combinations shared with PacBio from the Normal sample divided by the categories indicated in c) that mapped within the genomic location of the PacBio read (same genomic location as Pacbio) or in a different genomic location (different genomic location as Pacbio). The results were comparable across all samples. Results for the rest of the samples can be found in Supplementary Figure 2.

We found between 0.65% to 0.87% of the unique tag IDs to be mapped by PacBio data but unmapped in Illumina data. Those PacBio reads did not differ in GC content from the rest of the reads, they represented all isoform categories in similar proportions as the other read categories (Supplementary Fig 1). However, the average length of those reads was shorter than the average length of reads common between Illumina and PacBio (469-671 bp vs 713-856 bp) (Supplementary Fig 1).

The common cell barcode-UMI combinations found in both datasets were not equally counted to the gene matrix. PacBio retained a higher proportion of the shared tag IDs (43-64% vs 34-47%) and only 2-3% of them were multi-mapping, all of which were eliminated from final counts (Fig 2.c). Illumina data had a higher proportion of the shared fragments that were multi-mapping (12-16.6%) of which the majority (7-11% of all common tag IDs) were discarded from the final count matrix because of mapping to more than one gene. Of the uniquely mapping tag IDs in Illumina data (87-90% of the shared tag IDs), 95-98% of both the counted and not counted reads mapped within the same genomic range of the PacBio reads (Fig 2.d). On the other hand, from the multi-mapping Illumina tag IDs, 86-90% of those that were included in the UMI counting had at least one alignment within the same genomic location of the equivalent PacBio read but 97-98% of those that were discarded from the final count matrix did not align within the genomic location of the PacBio reads (Fig 2.d).

In order to better understand the reasons for the unique tag IDs in each dataset, we looked at the characteristics of shared and dataset-specific reads. In Illumina we found that the unique tag IDs had an elevated proportion of TSO and poly-A reads (Fig 3.a). The TSO removal step is part of the MAS-Seq library preparation protocol and thus such molecules are underrepresented in the PacBio reads (PacBio, 2023a). Between 67% and 84% of the Illumina reads mapped to exons, of which 30-59.4% were reads unique to Illumina counted to the gene count matrix (Fig 3.b). However, the fraction of reads unique to Illumina showed an overrepresentation of intergenic and intronic reads not counted into the final gene count matrix (25% of the Illumina-specific reads) (Fig 3.c). The intergenic reads were on average lower in GC content (44%), as expected from genome architecture (Piovesan et al., 2019), and the intronic reads showed an overrepresentation of reads from the >70% GC content category. The fraction of Illumina reads shared with PacBio were in majority exonic centred at 40-50% of GC content (Fig 3.d).

**Figure 3.**
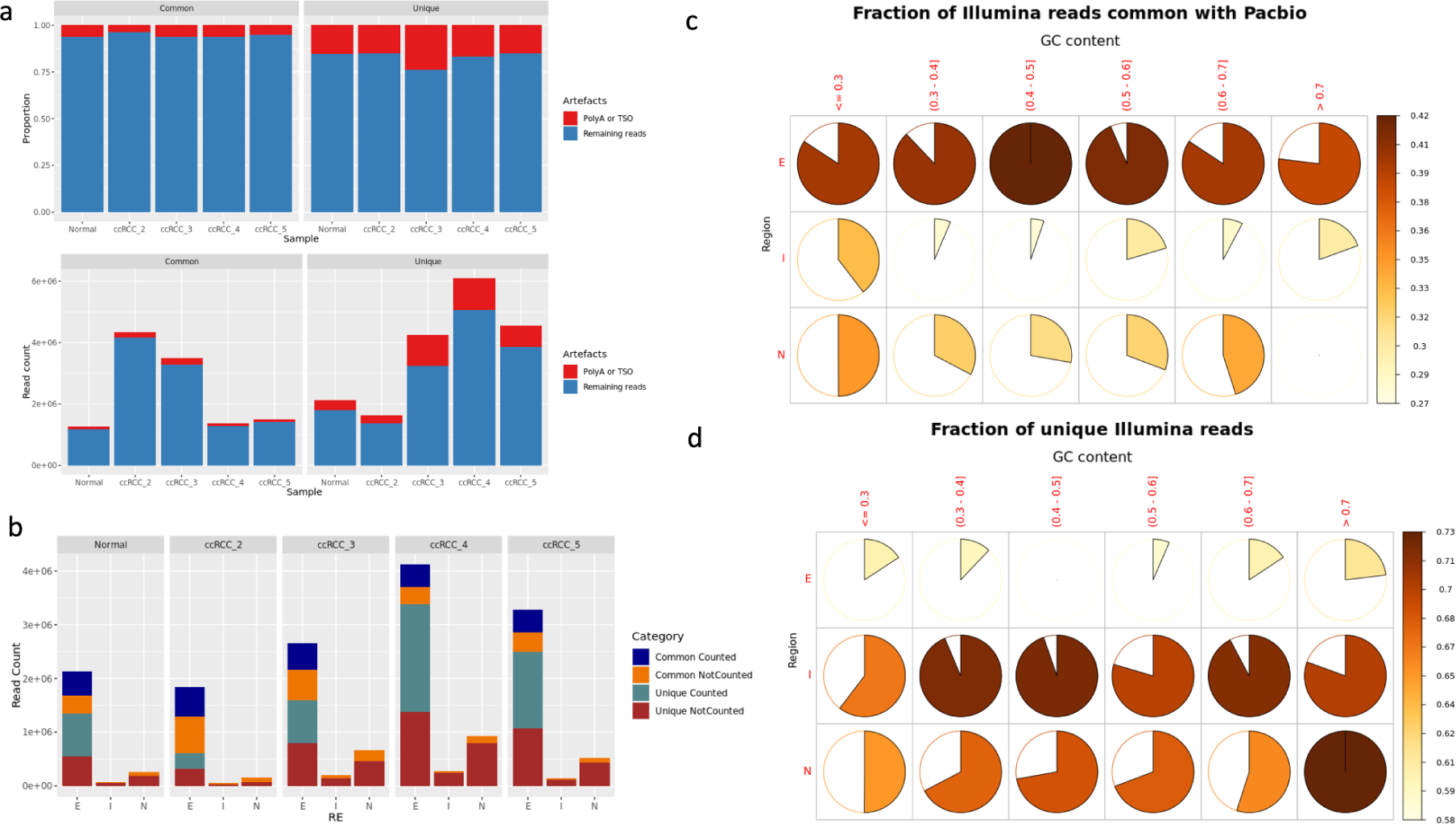
Characteristics of Illumina cell barcode-UMI combinations common with PacBio or unique to Illumina. a) The proportion of artefacts (polyA or TSO reads) in both categories. b) The number of reads mapping to exons (E), introns (N) and intergenic regions (I) and divided by whether they were counted in the final gene counts matrix. c) Fraction of Illumina reads shared with PacBio represented across different GC content categories (0.3-0.7) and different genomic regions. d). Fraction of reads unique to Illumina represented across different GC content categories (0.3-0.7) and different genomic regions.

The PacBio reads shared with Illumina showed an underrepresentation of reads shorter than 500 bp which can be explained by the size selection step that is part of the 10x Genomics 3’ library protocol for Illumina sequencing (Fig 4.a). Consequently, the reads unique to PacBio were on average shorter than the shared reads (487-679 bp vs 713-856 bp) (Fig 4.a). Between 75% and 84% of those reads were counted to the gene matrix and most of them mapped to isoforms labelled as fully overlapping with reference transcripts (Full Splice Match - FMS), or matching consecutive but not all splice junctions of the reference transcripts (Incomplete Splice Match - ISM) (Tardaguila et al., 2018) (Fig 4.b). The two isoform categories were represented in the same proportions in the reads common with Illumina.

**Figure 4.**
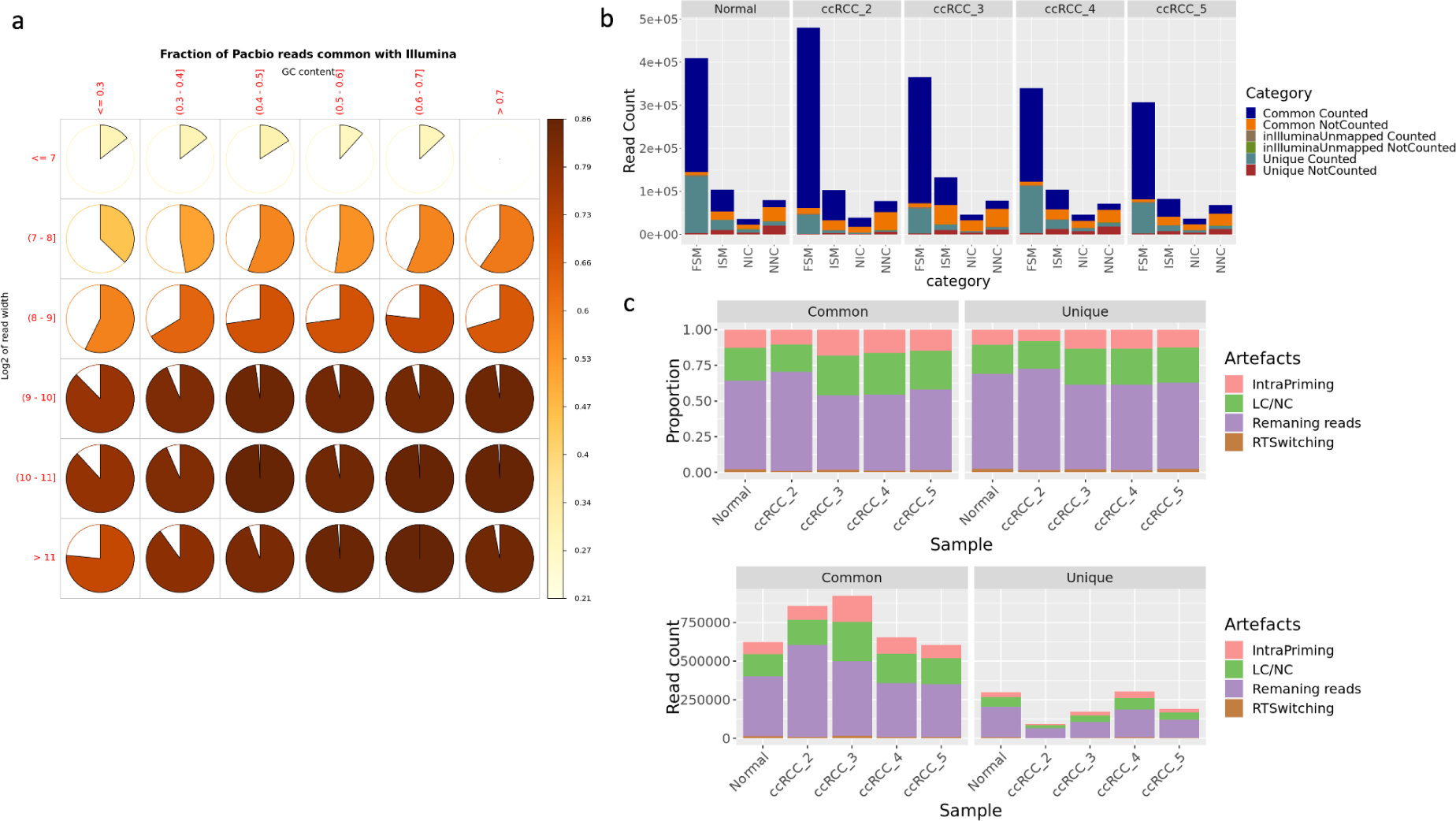
Characteristics of PacBio cell barcode-UMI combinations common with Illumina or unique to PacBio. a) Fraction of PacBio reads shared with Illumina represented across different GC content categories (0.3-0.7) and across different lengths (log2 of read length, 7-11). b) Number of reads mapping to isoforms from different categories including FSM - Full Splice Match (isoforms labelled as fully overlapping with reference transcripts), ISM - Incomplete Splice Match (isoforms matching consecutive but not all splice junctions of the reference transcripts), NNC - Novel Not in Catalog (isoforms harbouring at least one unknown splice site) and NIC - Novel In Catalog (isoforms presenting combinations of known splice donors and acceptors) (Dondi et al., 2023; Tardaguila et al., 2018). c) Proportion of intra-priming, reverse-transcriptase switching, low coverage/ non-canonical artefacts across the two groups of PacBio reads.

Long reads, due to providing isoform information, give an advantage of identifying additional data artefacts, including intra-priming or reverse-transcriptase switching transcripts (Tardaguila et al., 2018), on which we elaborate more below. Reads unique to PacBio had 1.5-7% fewer reads identified as mapping to the aforementioned artefacts than in the reads common with Illumina.

### Gene count matrix

Next, we evaluated the gene count matrix constructed from both datasets. For Illumina data, before the UMI count, CellRanger corrects UMI sequences and removes PCR duplicates. In our analysis, we additionally counted only reads mapping to exonic regions. For PacBio data, after retaining reads not mapping to any of the artefactual isoforms, we used *pigeon* to generate the gene-level count matrix. As previously mentioned, Illumina data consisted of more cells but we observed those cells to be of much lower quality than the cells found in both datasets. The cells had on average lower UMI count (Fig 1.b), fewer number of detected genes and on average elevated mitochondrial content (Fig 5.a). The median number of genes per cell in cells Illumina shared with PacBio was above 6300 across all samples whereas for those cells unique to Illumina, the median number of genes did not exceed 2650 across all samples (Fig 5.c). The median number of genes in PacBio data varied between 2672 and 5401 (Fig 5.c).

**Figure 5.**
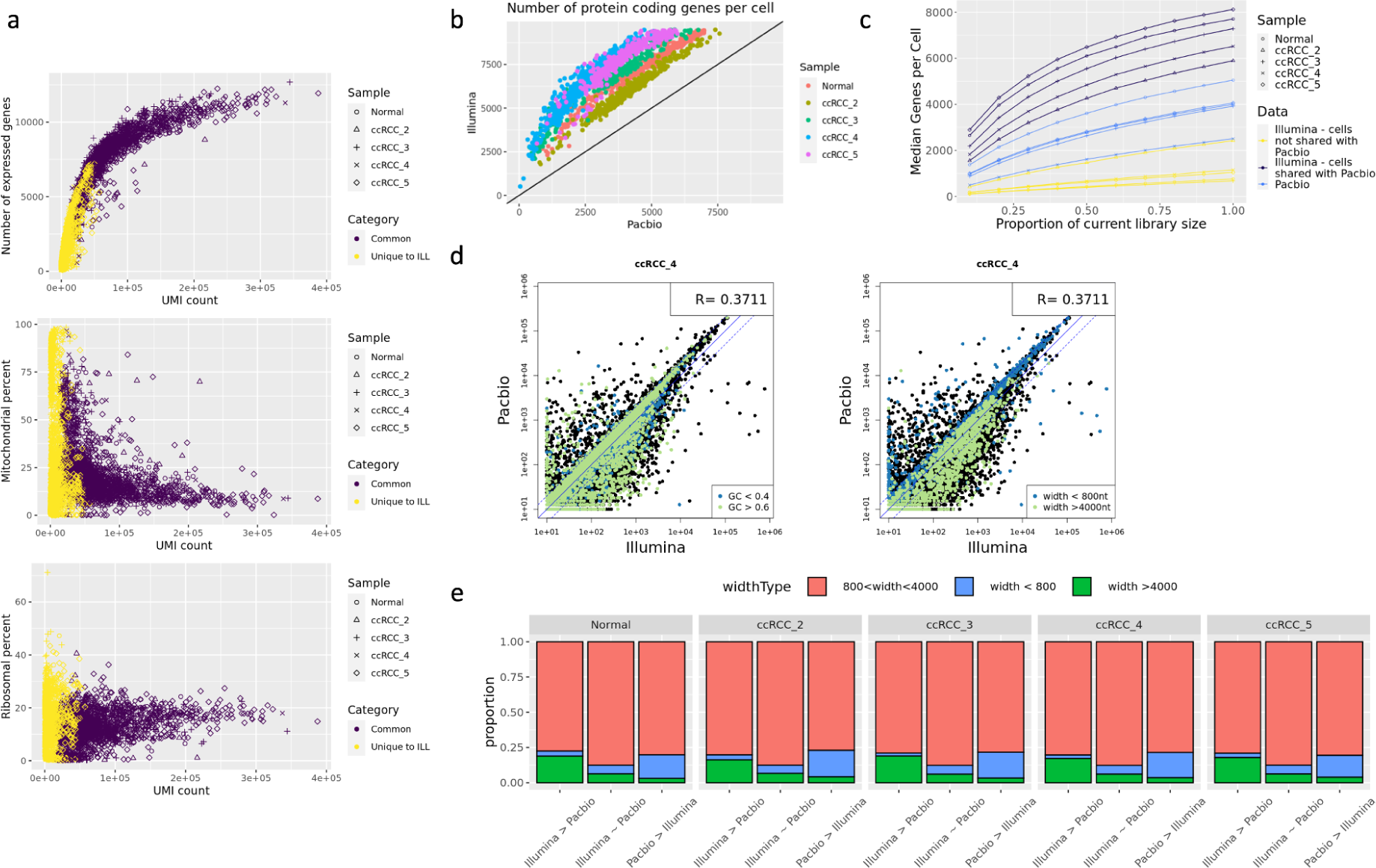
a) From top to bottom: number of expressed genes, fraction of reads mapping to mitochondrial genes and fraction of reads mapping to ribosomal genes correlated with UMI count per each cell in Illumina data categorised by whether the cell was also found in PacBio data. b) Correlation between PacBio and Illumina data in the number of protein coding genes per cell for same cells c) Median genes per cell correlated by the subsampled proportion of the library size in Illumina data separated by cells shared with PacBio and cells unique to Illumina and in PacBio data. d) Correlation between PacBio and Illumina data in the normalised sum of counts per gene across all common cells coloured by GC content and gene width (plots for remaining samples can be found in Supplementary Fig 5). e) Proportion of genes from three width categories (< 800 bp, between 800 bp and 4000 bp, and > 4000 bp) with significantly higher counts in Illumina (edgeR exactTest, logFC > 1, FDR < 0.05), with counts equal between Illumina and PacBio and with significantly higher counts in PacBio (edgeR exactTest, logFC < 1, FDR < 0.05).

For the subsequent comparison, we look only at cells detected in short- and long-read data. In short-read data, we detected a total of 15,219-17,017 protein coding genes associated with 19-52 million total UMIs, whereas in long-read data we detected 13,930-14,620 genes in a total of 5.7-13.9 million UMIs. Between 82-86% of the genes detected were shared between the two datasets (Supplementary Fig 3).

To observe the correspondence in counts for the same genes from the two datasets, we compared normalised and scaled sum of counts across all cells (pseudo-bulk). The counts correlated with a Pearson’s correlation coefficient varying between 0.37 and 0.59 (Fig 5.d). With edgeR exactTest we selected genes with significantly discordant counts including the genes missing from either of the datasets (logFC > 1, FDR < 0.05) and investigated their length and GC content. Genes with higher counts in Illumina (911-1629 genes) were found to be on average longer (2691-2819 bp) and have an overrepresentation of genes longer than 4000 bp (Fig 5.e). Genes with higher counts in PacBio (301-387 genes) were found to be on average shorter (1585-1672 bp) and consisted of a higher proportion of genes shorter than 800 bp (Fig 5e). The remaining genes with comparable counts between the two datasets had an average length of 1954 bp to 1984 bp. We did not find any difference in GC content between the three groups of genes (Supplementary Fig 4).

To understand if the genes with incongruent counts had a significant impact on the overall gene expression results, we merged the two datasets, projected the gene expression onto 2-dimensional embeddings using UMAP and measured the Euclidean distance between any two same cells. We found the datasets to closely overlap, with the Euclidean distance being lower than 2 for all but 1 or 2 cells. The exceptions were the Normal and the ccRCC_3 sample where 5 and 42 cells from both datasets had a distance equal or greater than 2, respectively. In sample ccRCC_3 the cells formed adjacent but distinct clusters (Fig 6.a), largely differentiated by the expression of mitochondrial genes (Fig 6.b and c). Indeed, across all samples, we found 0.1% to 74% difference in mitochondrial content between PacBio and Illumina for the same cells (Supplementary Figure 6).

**Figure 6.**
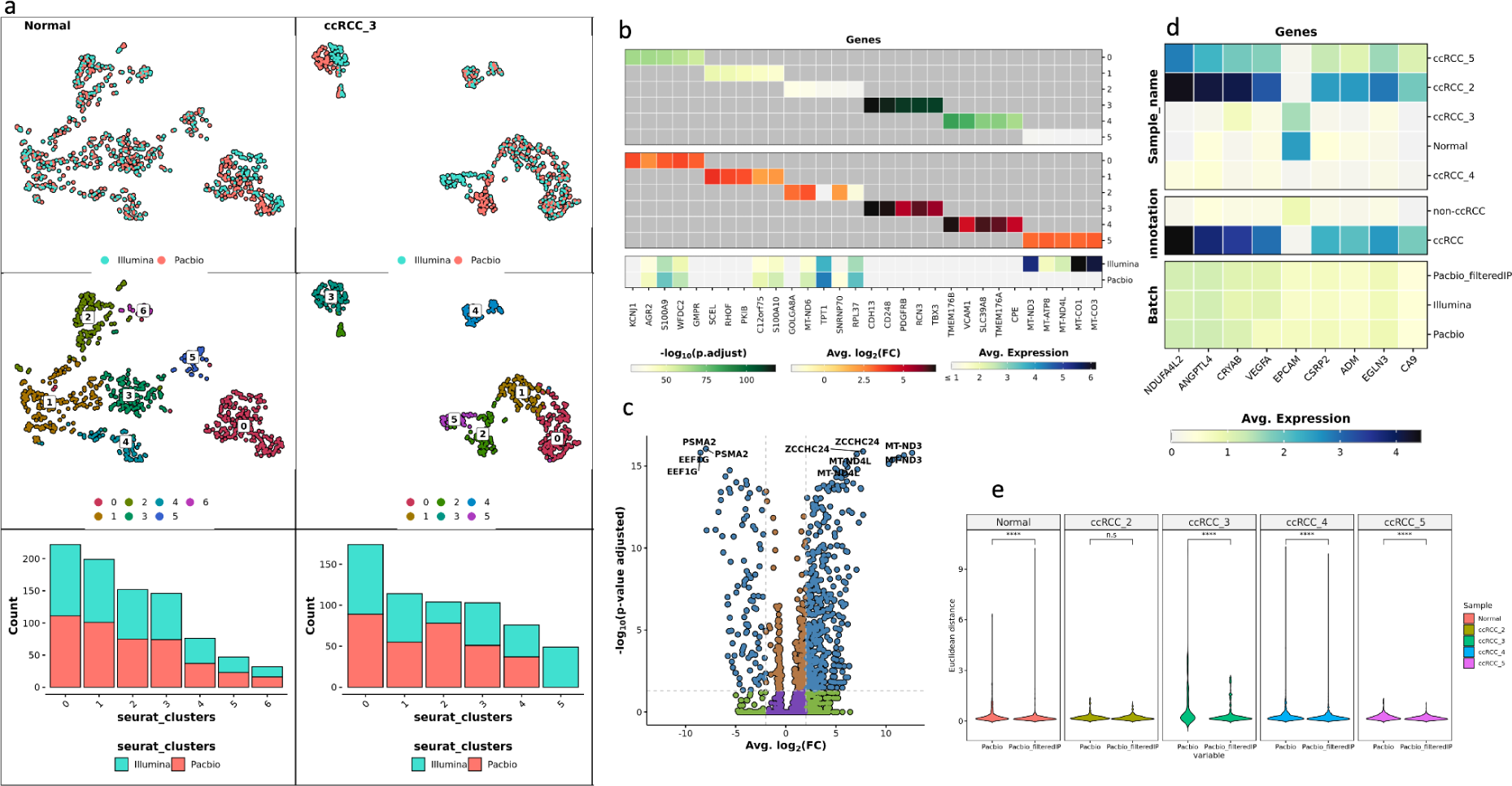
Clustering and gene expression comparison between Illumina and PacBio data. a) UMAP embeddings for the Normal and ccRCC_3 sample, coloured by sequencing method (top) or coloured by cluster identity (middle). Bottom: barplots representing the representation of each method in each cluster. The rest of the samples can be found in Supplementary Figure 7. b) Group-wise differential expression analysis results performed for sample ccRCC_3 between the different clusters. c) Differential expression analysis between PacBio and Illumina for cluster 3 for sample ccRCC_3. d) Gene expression of marker genes for ccRCC and epithelial cells divided by sample, cell annotation and data type. e) Comparison of the euclidean distance on UMAP plot between cells with the same barcodes from Illumina and 1) PacBio fully filtered data missing all artifactual isoforms (labelled Pacbio) and 2) PacBio data filtered only of intra-priming isoforms (labelled Pacbio_filteredIP). The significance of the difference in means was measured with a t-test.

The cell types were annotated according to CA9 expression as ccRCC and non-ccRCC (not malignant) cells, following guidelines from (Karakulak et al., n.d.). According to that maker, ccRCC_3 resembled the Normal sample and in ccRCC_2, ccRCC_4 and ccRCC_5 we detected 96.8%, 3.8% and 56% of cells expressing CA9, respectively. We looked at the expression of the CA9 gene as well as other markers characteristic for ccRCC pointed out by Karakulak et al. unpublished (Bi et al., 2021; Chen et al., 2023; Miikkulainen et al., 2019; Young et al., 2018; Zhang et al., 2021) and epithelial cells (Schutgens et al., 2019), expected to be present at high abundance in Normal sample, and found no difference in relative gene expression between PacBio data and Illumina data (Fig 6.d.).

### Long-read data filtering

PacBio reads are not only filtered from the TSO reads during library preparation but isoform information provides additional advantages for filtering out artefacts. The isoform filtering occurs in the *pigeon* workflow just before the construction of the final count matrix in the “Read Segmentation and Iso-Seq workflow” within SMRTLink v11.1 and distinguishes 3 types of artifactual transcripts: intra-priming (with adenine stretches in the genomic position downstream of the 3’ end), RT-switching (reverse transcriptase template switching) or low coverage/non-canonical (Tardaguila et al., 2018). Across all our samples between 39-54% of reads and between 79-87% of isoforms were flagged as belonging to any of these categories of artefacts and thus discarded from the final counts (Fig 7.a and 7.b). To investigate the contribution of those different artefacts on the gene counts, we compared the fully filtered data to data filtered only from intra-priming isoforms. Reverse transcriptase switching isoforms made up only 1.5-2.3% of the reads and 1.6-2.1% of the isoforms therefore we did not consider it to have a major contribution.

**Figure 7.**
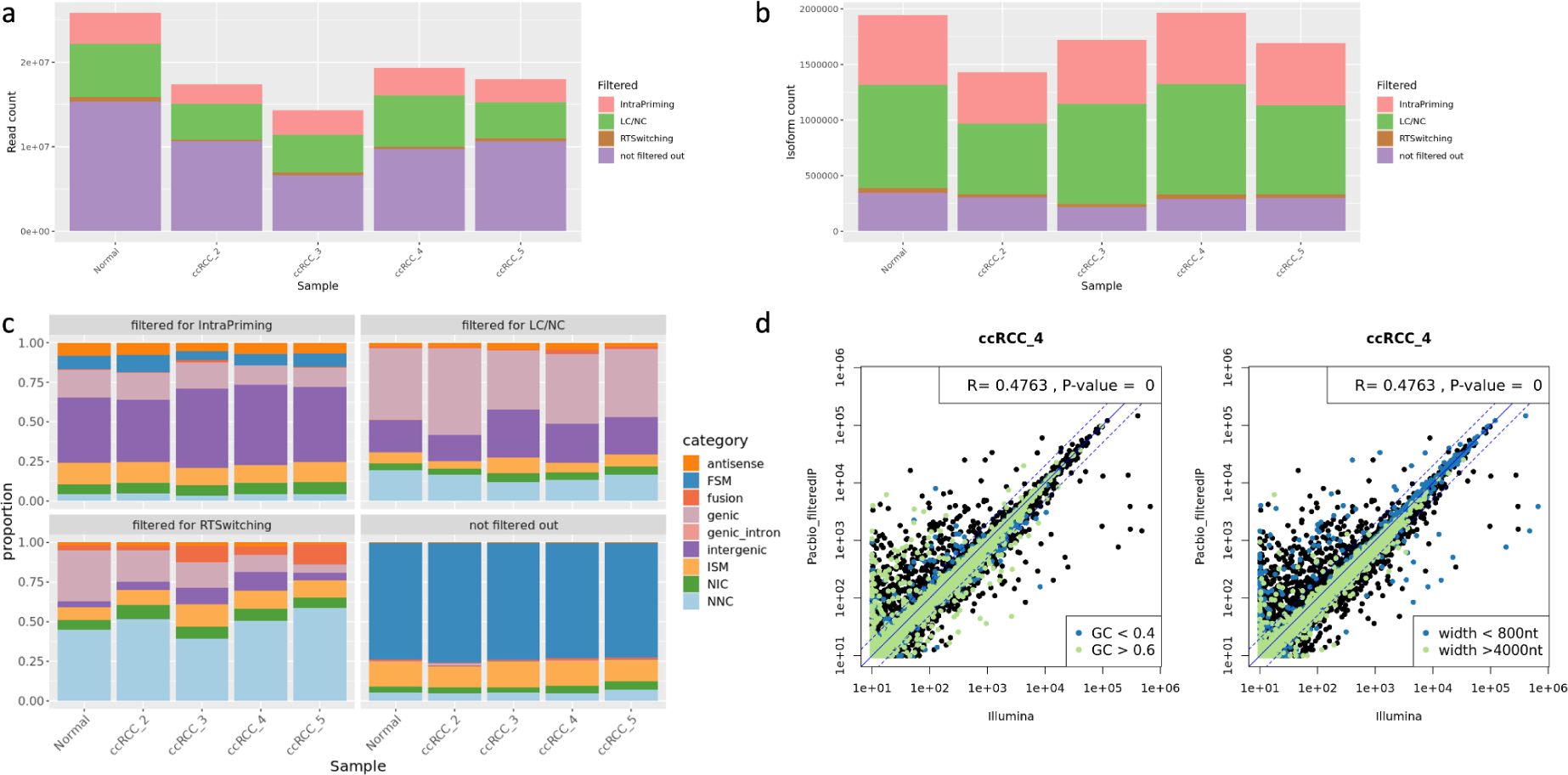
Characterization of PacBio data custom filtered from only intra-priming isoforms (and reads mapping to those isoforms). a) Number of reads filtered out due to mapping to intra-priming, low coverage/non-canonical (LC/NC) and RT switching isoforms and the reads remaining. b) Number of isoforms belonging to the four categories. c) Categorization of artefactual and remaining reads according to the isoform categories assigned by *pigeon* by sample. c) Correlation between Illumina and PacBio data filtered only of intra-priming isoforms in the sum of counts across all common cells coloured by GC content and gene width for sample ccRCC_4 (the plots for remaining samples can be found in Supplementary Fig 8).

The intra-priming transcripts constituted 32-34% of the isoforms and 13-20% of the reads. The reads were on average longer in length (1356-1529 bp vs 1323-1385 bp). According to *pigeon* classification, most of the reads were flagged as intergenic and intronic (39-51%) and genic (overlapping with introns and exons, so by definition, unspliced) (12-17%).

We found 44-52% of the isoforms and 24-31% of the reads to be assigned to the category of low coverage and non-canonical isoforms (Fig 7.a and 7.b). Majority of the reads were categorised as unspliced (genic, 37.5-54.4%), intergenic (16-30%) and novel (17.5-24%) (Fig 7.c). We took a sample of the Illumina reads with the same cell barcode-UMI combinations as the “genic” PacBio reads and also found 52-84% of them to not overlap with the splicing of the reference transcripts.

Retaining the low coverage/non-canonical reads in the analysis resulted in an increase in the total UMI count to 7.2 to 16.8 million which caused an increase in the mean and median UMI counts per cell by 18-27% and 30-51%, respectively, compared to the fully filtered data. However, we also observed an increase in mitochondrial content on average by 1.5-5.5%. The number of protein coding genes increased from 13,930 (ccRCC2) - 14,620 (ccRCC_5), as reported above, to 14,932 (ccRCC2) - 15,877(ccRCC_5). Between 88-90% of the genes detected were shared with the Illumina data. Not only did we observe a rise in the total number of genes but also a change in gene counts - the correlation coefficient between the normalised sum of counts across all cells between PacBio data filtered of only intrapriming transcripts and Illumina data increased to 0.48-0.73. With edgeR exactTest we again selected genes with significantly discordant counts including the genes missing from either of the datasets (logFC > 1, FDR < 0.05) and this time found only 112-211 genes with higher counts in Illumina data, which constituted only about 12% of the genes found in comparison of the fully filtered PacBio data. The average length of those genes did not show any difference from the genes with equal counts between the two datasets (1632-2057 bp vs 2032-2058 bp). We found 461-602 genes with higher counts in PacBio data, which was slightly higher than detected previously. Genes with significantly higher counts in PacBio data were again on average shorter (1755-1826 bp) and exhibited an overrepresentation of genes below 800 bp long (Supplementary Fig 9).

We merged the PacBio data filtered only of the intra-priming isoforms and with the Illumina data and projected the gene expression of the three datasets onto a UMAP. With the increase in the number of genes and UMI counts per cell, we observed a significant decrease by 23-50.5% in the mean Euclidean distance between the same cells in Pacbio and Illumina data for four out of the five samples in comparison to the distance measured in the clustering analysis of fully filtered Pacbio data with Illumina (Fig 6.e). Except for ccRCC_3, we did not observe any changes in clustering results (Supplementary Figure 10) or in expression of the ccRCC specific markers (Fig 6.d). In clustering of ccRCC_3 we observed the previously mentioned adjacent clusters to now overlap (Supplementary Figure 10).

## Discussion

Long-read sequencing applied to scRNA-seq allows for deciphering of cell heterogeneity based on full-length transcripts providing variant and isoform information. Recent developments have expanded their use (Pardo-Palacios et al., 2023) and allowed their decoupling from short reads but recent evidence still suggests a certain degree of variability in information obtained from long reads when compared to short reads (Dondi et al., 2023). Here we used single cells from four samples of ccRCC-derived organoid and one matched healthy kidney organoid sample to answer the question of the extent of comparability between the long- and short-read data and the reason for the differences. We focused on both data processing and output at two steps: mapping of reads to the reference genome and gene-wise UMI counting.

We observe a great overlap in the information provided by both datasets. We find the same set of high quality cells. High proportion of cell barcode-UMI combinations are recovered by both methods. Gene expression results per cell connect two same cells in non-linear dimensionality reduction analyses except for cells with high expression of mitochondrial genes observed in Illumina data. Cell markers specific to ccRCC show comparable normalised expression. We do, however, observe some variability due to variation at different stages of the protocols and due to the nature of the two sequencing methods.

Short-read sequencing data still provides higher coverage at a lower cost. Following the guidelines for sequencing depth from PacBio for Sequel IIe, we still recover less UMIs per cell and fewer expressed genes. However, a higher proportion of Illumina-specific reads were not counted due to mapping to introns and intergenic regions or mapping to multiple locations in the genome. Long-read information allowed for more precise allocation of same reads to unique genomic regions.

Paradoxically, sequencing full-length transcripts with long reads allows recovery of short transcripts (<125bp) which are otherwise discarded in Illumina 10x library protocol due to transcript fragmentation and size selection steps. Removing short fragments by size selection leads to underestimation of counts for short genes (< 800 bp), also observed by Dondi et al (2023). Additionally, the MAS-seq library protocol also includes TSO artefact removal which is advantageous for saving sequencing efforts. In cell barcode-UMI combinations unique to Illumina we identified a higher proportion of TSO and PolyA artefacts.

Isoform information from long reads provides more information on the types of sequencing artefacts and thus leads to more precise filtering of reads for gene expression profiling. We identified 33.5-36.3% of isoforms and up to 22% of reads to be discarded due to being flagged as intra-priming or reverse-transcriptase switching. We observed the low coverage/non-canonical isoforms to have the biggest impact on the UMI counting causing a decrease in counts especially for genes longer than 4000 bp. We identified up to 52% of isoforms and up to 31% of reads to belong to that category. The percentage was higher than identified by Dondi et al. (Dondi et al., 2023) which might be caused by the fact their sequencing was done at a higher depth. However, the largest proportion of those reads was identified as unspliced (containing introns and exons) which suggests an impact of sample specific cellular state (Riba et al., 2022). A large proportion of those reads were also identified as unspliced in Illumina data. Discarding them from counting would thus follow the logic that the library protocol is aimed to target mature transcripts. However, up to 24% were identified as a novel and, as revealed by the LRGASP consortium, identification and quantification of lowly expressed and complex transcripts continue to be a challenge in long reads (Pardo-Palacios et al., 2023). Experimental validation of those transcripts would provide information on whether they are biologically significant or true artefacts. Non-canonical transcripts have been shown to encode tumour-specific antigens therefore it would be beneficial to verify the usefulness of this filtering step in the workflow and whether the filtering needs to be done in a sample-specific manner (Apavaloaei et al., 2020; Camarena et al., 2023). The significance and frequency of novel isoforms from the unfiltered, remaining reads is addressed in (Karakulak et al., n.d.).

Our aim was to test the most frequently applied workflows on both datasets and we do acknowledge the limitations, for example in comparing bam files resulting from two different mapping algorithms. We also sequence a lower number of cells than is perhaps of interest in most scRNAseq experiments. However, we think that our benchmark analysis identified multiple sources of variations between long- and short-read sequencing technologies. The finding is informative and has a general applicability. With increasing throughput in long-read sequencing, we would expect even a greater overlap in the obtained data and thus we think long reads provide an alternative to short-read sequencing for single-cell experiments.

## Methods

### Generation and characterization of ccRCC Patient-Derived Organoid Samples

The Department of Pathology and Molecular Pathology at University Hospital Zürich made available the patient tissue samples. They were collected and biobanked according to previously described procedures (Bolck et al., 2019). The study was approved by the local Ethics Committee (BASEC# 201 9-01 959) and in agreement with Swiss law (Swiss Human Research Act). All patients gave written consent. Organoids were established as previously described (Bolck et al., 2021). The details of the methods are described in (Karakulak et al., n.d.).

### Library preparation

#### 10x Genomics library preparation and short-read sequencing

The samples were analysed first using the 10x Genomics Chromium platform (Zheng et al., 2017). Library preparation was conducted following the 10x Genomics Chromium Single Cell 3’ Reagent Kits User Guide (v3.1 Chemistry Dual Index). Organoids were dissociated, washed to eliminate debris, clumps, and contaminants, and resuspended in a 1x PBS/0.04% BSA solution at a concentration of 500 cells/µL. Cell viability and concentration were determined using a LUNA-FX7 Automated Cell Counter (Logos). To ensure a greater sequencing depth per cell using the PacBio platform, our target was to recover approximately 700 cells per library preparation. The cells were then combined with a master mix containing reverse transcription reagents. The single-cell 3’ v3.1 gel beads, which carry the Illumina TruSeq Read1, a 16bp 10x barcode, a 12bp UMI, and a poly-dT primer, were loaded onto the chip along with oil for the emulsion reaction. The Chromium X partitioned the cells into nanoliter-scale gel beads in emulsion (GEMs), where reverse transcription occurred. All cDNAs within a GEM, representing one cell, shared a common barcode. After the reverse transcription reaction, the GEMs were broken, and the full-length cDNAs were captured by MyOne SILANE Dynabeads and then amplified. The amplified cDNA underwent cleanup with SPRI beads, followed by qualitative and quantitative analysis using an Agilent 4200 TapeStation High Sensitivity D5000 ScreenTape and Qubit 1X dsDNA High Sensitivity Kit (Thermo Fisher Scientific). The cDNA was enzymatically sheared to a target size of 200-300 bp, and Illumina sequencing libraries were constructed. This process included end repair and A-tailing, adapter ligation, a sample index PCR, and SPRI bead clean-ups with double-sided size selection. The sample index PCR added a unique dual index for sample multiplexing during sequencing. The final libraries contained P5 and P7 primers used in Illumina bridge amplification. Sequencing was performed using paired-end 28-91 bp sequencing on an Illumina Novaseq 6000 to achieve approximately 300,000 reads per cell.

#### MAS-Seq library preparation and long-read sequencing

The single-cell full-length cDNA generated using the 10x Genomics Chromium Single Cell 3’ Reagent Kits (v3.1), as described above, with the input amount of 45 ng/sample, was directed for single-cell MAS-Seq (Multiplexed Arrays Sequencing) libraries preparation using the MAS-Seq for 10x Single Cell 3’ kit (Pacific Bioscience, CA, USA). Firstly, template switch oligo (TSO) priming artefacts generated during 10x cDNA synthesis were removed in the PCR step with a modified PCR primer (MAS capture primer Fwd) to incorporate a biotin tag into desired cDNA products followed by their capture with streptavidin-coated MAS beads. cDNA free from TSO artefacts was further directed for the incorporation of programmable segmentation adapter sequences in 16 parallel PCR reactions/sample followed by directional assembly of amplified cDNA segments into a linear array. Such formed MAS arrays with the average length of 10-15 kbp were further DNA damage repaired and nuclease treated in order to produce final single-cell MAS-Seq libraries which quantity and quality were measured by Qubit 1X dsDNA High Sensitivity Kit (Thermo Fisher Scientific) and pulse-filed capillary electrophoresis system Femto Pulse (Agilent), respectively. Each single-cell MAS-seq library was used to prepare the sequencing DNA-Polymerase complex using 3.2 binding chemistry (Pacific Bioscience) and further sequenced on a single 8M SMRT cell (Pacific Bioscience), on Seque IIe sequencer (Pacific Bioscience) yielding in ∼ 2 M HiFi reads and∼ 30M segmented reads/ sample.

### Data analysis

#### Short-read sequencing

For direct comparison with PacBio data, we generated a CellRanger compatible reference for the human genome version GRCh38.p13 and GENCODE annotation v39 using the files available via SMRTLink v11.1. The raw data was mapped to this reference and filtered with CellRanger v7.2.0 (Zheng et al., 2017). Bam files were subsampled for cell barcodes starting with *AA* sequence, for handling feasibility, and analysed using the GenomicAlignments R package v1.38.0 (Lawrence et al., 2013). For those reads where the UB tag (corrected UMI) was missing, we substributed with the UR tag (UMI tag reported by the sequencer) and the data was filtered from PCR duplicates. Intronic reads were excluded from UMI counting. The filtered feature counts were converted to a Seurat object and filtered for cells common with PacBio data and for protein coding genes. Ribosomal and mitochondrial content was calculated as the proportion of counts assigned to ribosomal or mitochondrial genes, respectively. Cells common with PacBio were annotated as ccRCC or non-ccRCC (healthy) according to the annotation done by (Karakulak et al., n.d.). Pseudo-bulk was generated by obtaining the sum of counts for each gene across all cells. Expression data was then analysed using the seurat workflow (v5.0.1) (Hao et al., 2021, 2023): the counts were normalised using log normalisation with a scale factor of 10,000 and subsequently scaled. We selected 3000 highly variable features using the vst method. Linear dimensional reduction was used to determine dimensionality of the dataset and top 30 PCs were used for non-linear dimensionality reduction analyses (UMAP/tSNE). Ambient mRNA estimation was performed with SoupX (1.6.2) (Young & Behjati, 2020). For analysis of whether reads annotated as “genic” (containing exons and introns) in PacBio data are also detected as unspliced in Illumina data, we subsampled 1500-2000 cell barcode-UMI combinations of the “genic” set from CellRanger bam files and checked whether they were compatible with the splicing of the reference transcripts (using GenomicAlignments R package).

#### Long-read sequencing

Long-read sequencing data was processed using the “Read Segmentation and Iso-Seq workflow” within SMRTLink v11.1. For two samples (Normal and ccRCC_2), for each of which three SMRTcells were sequenced, all the sequencing data was merged before analysis to increase data coverage. No confounding batch effects were observed before merging. Briefly, within the pipeline, HiFi reads (high fidelity CCS reads with QV > 20) were deconcatenated into segmented reads based on segmented adapters with *skera* tool. The reads were then processed with *isoseq* (v3.8.1 within SMRTLink v11.1) for removal of cDNA primers and barcode and UMI tags, reorientation, trimming of poly-A tails, cell barcode correction, real cell identification and PCR deduplication via clustering by UMI and cell barcodes. Reads were then aligned to the GRCh38.p13 genome with *pbmm2 align* (a wrapper around minimap2). Bam files were analysed using the GenomicAlignments R package, selecting cell barcodes matching the subsampled Illumina cell barcodes (Lawrence et al., 2013). PacBio cell and UMI barcodes are written in reverse complement and this has been accounted for in our analyses. Unique isoforms were collapsed and classified with *pigeon*. Filtering out isoforms was done in two ways: following the default parameters of filtering intra-priming, RT-switching and low-coverage/ non-canonical isoforms, and only filtering the intra-priming isoforms. For both datasets a gene count matrix was created with *pigeon make-seurat*. Results from both filtering steps were compared for better understanding of the effect of the artefacts on gene counts. Both gene count matrices from PacBio were then subsequently used for comparison with Illumina data subsetted to only protein coding genes and common cells. Cells were annotated as ccRCC or non-ccRCC (non-malignant) with scGate (v1.6.0) using CA9 as a positive expression marker for ccRCC cells (Andreatta et al., 2022). The annotation was done by (Karakulak et al., n.d.). Ribosomal and mitochondrial content was calculated as the proportion of counts assigned to ribosomal or mitochondrial genes, respectively. Pseudo-bulk was generated by obtaining the sum of counts for each gene across all cells. The data was processed with seurat following exactly the same workflow as described above for Illumina.

#### Comparison

The long and short-read datasets were compared at the level of: 1) transcribed molecules and their alignments, and 2) gene count matrices. Since the protocol generates cDNA molecules tagged with a cell barcode and a UMI, we can directly match and compare the short- and long-reads generated from the same original molecule. We define for each molecule the tag id as the combination of cell barcode and UMI, and use this tag id to identify corresponding reads across the technologies. This allows us to identify molecules preferentially detected by each technology. Additionally, we evaluate characteristics like genomic location of alignments, proportion of artefacts, proportion of reads discarded from the final counts. The final gene count matrices were first analysed independently and compared without subsetting. Pearson’s correlation coefficient between Illumina and PacBio pseudo-bulk was calculated using stats R package (4.3.2). EdgeR exactTest (from edgeR 4.0.1) with dispersion coefficient of 0.1 was used to select genes with higher counts in Illumina or higher counts in PacBio including genes missing from either of the datasets. The p-values were corrected with the p.adjust function from EdgeR. The length and GC content of each gene was assumed to be the average length and GC content of all the reference isoforms. Subsequently, for cell-to-cell comparison, Illumina and PacBio seurat objects for each sample were merged using only common cells. Each data was processed independently (normalised, scaled and subjected to selection of 3000 highly variable features) before merging and the dimensionality reduction analyses were repeated after merging. The data was visualised using seurat (v5.0.1) and SCpubr (v2.0.0). Euclidean distance was measured between any two same cells from PacBio and Illumina on a UMAP. Because Illumina data was compared to two PacBio datasets generated with different isoform filtering options, a t-test using ggpubr (v0.6.0) was performed to compare the difference in mean euclidean distance per sample.

## Data Access

The code used for data analysis is available on github:

https://github.com/zajacn/scRNAseq_Long_reads_vs_Short_reads

The raw data has been deposited in XXX under XXX accession.

## Acknowledgments

We would like to thank Mark Robinson for valuable discussion during the analysis process. We thank Krebsliga Zürich and EMDO Stiftung for funding.

## Supplementary Figures

Supplementary Figure 1. a. Analysis of cell barcode-UMI combinations that were mapped to the human reference genome from the PacBio data but were unmapped by Illumina. A. Read length distribution B. GC content distribution C. Proportion of reads belonging to different isoform categories.

Supplementary Figure 2. The proportion of Illumina cell barcode-UMI combinations common with Pacbio, divided by whether they were uniquely or multimapping and whether they were counted into the gene count matrix or not, that fall within the genomic range of the PacBio reads with the same cell barcode -UMI combination (TRUE, at least one alignment) and that do not fall within the genomic range of the Pacbio read (FALSE, none of the alignments).

Supplementary Fig 3. A. Distribution of UMIs per cell B. Distribution of total genes per cell. C. Overlap in detected genes between Illumina and PacBio.

Supplementary Figure 4. Proportion of genes with average (0.4-0.6), low (<0.4) and high (>0.6) GC content in genes with comparable counts between Illumina and PacBio (Illumina ∼ PacBio), with counts significantly higher in Illumina than in PacBio (Illumina > Pacbio) and vice versa (Pacbio > Illumina).

Supplementary Figure 5. Correlation between PacBio and Illumina data in normalised sum of counts per gene across all common cells for samples: Normal (healthy), ccRCC_2, ccRCC_3 and ccRCC_5.

Supplementary Figure 6. Correlation between Illumina and PacBio in mitochondrial content (in percentage) for the same cells.

Supplementary Figure 7. UMAP clustering analysis results for each sample on merged Illumina data with PacBio data filtered from all artefacts.

Supplementary Figure 8. Correlation between PacBio (filtered only from intra-priming isoforms) and Illumina data in normalised sum of counts per gene across all common cells for samples Normal (healthy), ccRCC_2, ccRCC_3 and ccRCC_5.

Supplementary Figure 9. Proportion of genes from three width categories (< 800 bp, between 800 bp and 4000 bp, and > 4000 bp) with significantly higher counts in Illumina (edgeR exactTest, logFC > 1, FDR < 0.05), with counts equal between Illumina and PacBio and with significantly higher counts in PacBio, using PacBio data filtered only from intra-priming isoforms.

Supplementary Figure 10. UMAP clustering analysis results for each sample on merged Illumina data and PacBio data filtered of only intra-priming isoforms.

## Supplementary Tables

Supplementary Table 1. Long-read (PacBio) and short-read (Illumina) raw data summary.

Supplementary Table 2. Summary of the subsampled data for analysis of bam files.

